# AI-Guided CRISPR Screen Accelerates Discovery of New Drug Targets

**DOI:** 10.64898/2026.02.26.708368

**Authors:** Chenlin Zhao, Mushaine Shih, Sharif Ahmed, Steven Song, Amber Lennon, Julia M Mayes, Mengqi Jonathan Fan, Pei-Ying Lo, Jason Perera, Alanur Tutar, Andrew Kang, Elizabeth Warren, Ryan McClure, Aly A Khan, Shana O Kelley, Abdalla M Abdrabou

## Abstract

Psoriasis affects over 125 million people worldwide, yet the mechanistic understanding of keratinocyte-driven inflammation remains incomplete, limiting therapeutic innovation beyond costly systemic biologics that are prone to side effects. Here, we performed the first genome-wide CRISPR knockout screen in primary human adult epidermal keratinocytes to systematically identify regulators of IL-17 receptor A (IL17RA), a central node in psoriatic inflammation. To prioritize therapeutically tractable targets from over 19,000 screened genes, we integrated a large language model – *VirtualCRISPR* – trained on functional genomics data, identifying arachidonate 5-lipoxygenase (ALOX5) and oxytocin receptor (OXTR) as high-confidence novel hits with minimal prior association with psoriasis. Multi-omics validation revealed that ALOX5 and OXTR regulate IL17RA expression through distinct signaling pathways – ALOX5 through lipid mediators that stabilize the receptor at the cell surface, and OXTR through calcium signaling that reprograms cellular metabolism. Topical delivery of their inhibitors Zileuton (ALOX5) and Cligosiban (OXTR) exhibited therapeutic efficacy comparable to systemic anti-IL17RA antibody in the imiquimod-induced psoriasis model, suppressing pathogenic Th17/Tc17 responses, polarizing macrophages toward anti-inflammatory phenotypes, and normalizing epidermal hyperproliferation. Proteomic profiling in human 3D organotypic skin and murine models confirmed on-target pharmacology and revealed convergent suppression of neutrophil-keratinocyte inflammatory circuits. The use of VirtualCRISPR significantly shortened the timescale from screen to the identification of druggable hits with robust validation, and this work establishes a blueprint for integrating AI-driven target prioritization with functional genomics to accelerate therapeutic discovery.

## Introduction

Psoriasis is a chronic inflammatory skin disorder affecting 2-3% of the global population, characterized by keratinocyte hyperproliferation and immune dysregulation^1,2^. The IL-17/IL-17RA axis represents a critical pathway in psoriasis pathogenesis, and biologics targeting this axis (secukinumab, ixekizumab, brodalumab) have demonstrated remarkable clinical efficacy^3,4^. However, the cellular mechanisms governing IL-17 signaling in keratinocytes, the primary effector cells in psoriatic lesions remain incompletely understood.

While biologics have revolutionized psoriasis treatment, their requirement for systemic administration, high cost, and potential immunogenicity limit long-term use^5^. Topical corticosteroids, the mainstay of localized therapy, cause tachyphylaxis and skin atrophy with chronic use^6^. These constraints underscore an urgent need for novel small-molecule therapeutics amenable to topical delivery.

Functional genomics approaches, particularly CRISPR-Cas9 loss– and gain-of-function screens, have transformed our ability to dissect disease mechanisms and systematically identify therapeutic targets^7,8^. However, despite the central role of keratinocytes in psoriasis, genome-wide CRISPR screens in primary human keratinocytes remain technically challenging due to limited proliferative capacity and low transduction efficiency^9^. Consequently, although recent studies have begun to employ primary human keratinocytes for genetic screens, a comprehensive genome-wide CRISPR knockout screen has not been reported, potentially missing crucial keratinocyte-specific regulatory mechanisms^10,11^.

Here, we performed the first genome-wide CRISPR knockout screen in primary human adult epidermal keratinocytes (HEKa cells), identifying regulators of IL-17 receptor A (IL17RA) surface expression (Fig. 1A). To navigate the complex landscape of screening hits and extract biologically meaningful signals, we employed a large language model-based framework to prioritize candidates by novelty and causality (Fig. 1B, C)^12^, then identified small molecule inhibitors and FDA-approved drugs that could be repurposed to target the hits.

**Fig. 1.**
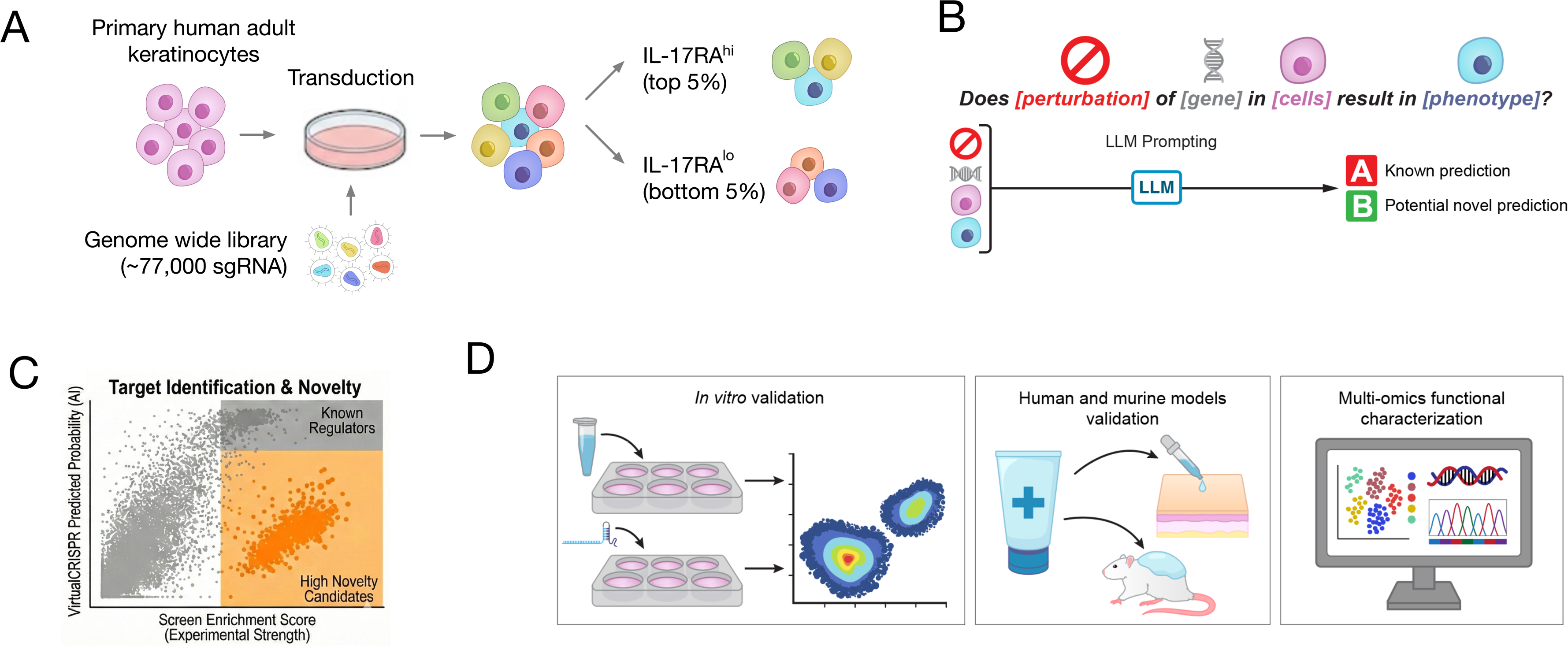
| Integrated CRISPR-Cas9 screening and AI-guided prioritization platform identifies IL17RA regulators in primary human keratinocytes. A, Experimental workflow. Primary human adult epidermal keratinocytes (HEKa) were transduced with the genome-wide Brunello sgRNA library (∼77,000 sgRNAs targeting ∼19,000 genes), selected with puromycin, expanded for five population doublings, and sorted by fluorescence-activated cell sorting (FACS) to isolate IL17RA^hi^ and IL17RA^lo^ populations (top and bottom 5%). Genomic DNA was extracted and sgRNA representation was determined by next-generation sequencing. B, AI-based hit prioritization strategy. The VirtualCRISPR model, trained on the GPT embeddings and BioGRID-ORCS database, predicted hit probability for each screened gene by addressing the question: “Does {perturbation} of {gene} in {cell type} result in {phenotype}?” Outputs were classified as known predictions (high model-predicted probability) or potential novel predictions (low model-predicted probability). C, Target identification and novelty classification. Enriched screen hits were plotted by experimental enrichment score (x-axis) versus VirtualCRISPR predicted probability (y-axis). Genes with high experimental enrichment and high predicted probability correspond to known regulators (gray region); genes with high experimental enrichment and low predicted probability represent high-novelty candidates with uncharacterized biology (orange region). D, Validation pipeline. Top novel candidates underwent targeted CRISPR-Cas9 knockout in primary keratinocytes, followed by multi-omics profiling and functional validation in three-dimensional organotypic human skin cultures and the imiquimod-induced mouse model of psoriasiform dermatitis.

Subsequently, we validated the hits through rigorous multi-omics analysis in vitro and in vivo in human and murine models (Fig. 1D). Topical delivery of Cligosiban (an OXTR antagonist) and Zileuton (an ALOX5 inhibitor) achieved efficacy equivalent to systemic anti-IL17RA antibody in the imiquimod-induced psoriasis model, linking neurohormonal and lipid signaling to IL-17-driven inflammation.

Furthermore, we demonstrate that integration of AI-powered target prioritization with genome-wide functional genomics accelerates therapeutic discovery. The VirtualCRISPR model, trained on thousands of CRISPR screens across diverse biological contexts, was applied not as a hit ranker but as a novelty filter: genes with strong experimental enrichment and high model-predicted probability reflect established biology, whereas genes with strong experimental enrichment and low predicted probability represent mechanisms not previously linked to the queried phenotype. OXTR and ALOX5 were identified as high-novelty candidates on this basis, each exhibiting predicted probabilities lower than IL17RA knockout itself despite robust experimental enrichment, indicating their roles in IL17RA regulation in keratinocytes were not anticipated from prior functional genomic knowledge.

Subsequent validation confirmed both as druggable nodes that recapitulate IL-17 receptor blockade efficacy when inhibited topically. This integrated platform, combining primary cell functional genomics, large language model-based novelty prioritization, and multi-omics validation, provides a generalizable blueprint for rational drug discovery in complex inflammatory diseases.

## Results

To systematically identify regulators of IL-17 receptor A (IL17RA) expression in HEKa, we developed a FACS-based CRISPR screen enabling isolation of two distinct cellular populations representing the top and bottom ∼5% of surface IL17RA-expressing cells. The stringent gating strategy ensured the accurate capture of both true positive and negative regulators of IL17RA expression. Preliminary validation using wild-type (WT) and IL17RA knockout cells confirmed the ability of the sorting to reproducibly separate WT and low surface IL17RA-expressing populations (Supplementary Fig. 1).

HEKa cells are often refractory to genetic manipulation, posing substantial challenges for large-scale perturbation screens^13^. To address this, we conducted preliminary experiments optimizing transduction efficiency while maintaining cell viability and library representation. Transduction enhancers such as polybrene markedly inhibited HEKa proliferation even at low concentrations (Supplementary Fig. 2A), excluding them from all transduction protocols. Although some chemical modulators like ROCK1 inhibitors can overcome the growth-shunting effect, they may introduce pathway-specific biases. Instead, we adopted a spinoculation-based transduction method in the absence of polybrene, followed by a 24-hour viral incubation period, achieving optimal library delivery at low MOI (∼0.25-0.35) (Supplementary Fig. 2B-D).

Using these optimized conditions, we transduced HEKa cells of 2 different healthy human donors with the human Brunello genome-wide CRISPR knockout library at a multiplicity of infection (MOI) of ∼0.35, ensuring single-guide representation per cell. Following puromycin selection, the transduced pool was expanded for five population doublings to allow for effective gene editing and depletion of essential genes. We then FACS-sorted the library-transduced population to isolate the top and bottom 5% of IL17RA surface expressing cells (Fig. 1A).

Each donor yielded approximately 80 million keratinocytes before sorting. We extracted genomic DNA from IL17RA^high^ and IL17RA^low^ populations and performed deep sequencing to quantify guide RNA enrichment in each sorted pool. Sorting produced a clear shift in IL17RA expression, confirming robust phenotypic separation (Supplementary Fig. 3). To evaluate library representation, ∼25 million cells were cryopreserved immediately after puromycin selection (T₀).

We applied MAGeCK analysis pipeline to compute FDR and log₂ fold-change values and identify genes whose knockout altered IL17RA expression. We retained more than 98% of the sgRNAs at the T_0_ population, indicating a high library coverage after transduction (Supplementary Fig. 4A). Using an FDR threshold of 0.1 and requiring at least two sgRNAs per gene, we identified significantly enriched genes in IL17RA^low^ populations (Supplementary Fig. 4B).

As expected, IL17RA sgRNAs were among the top hits enriched in the IL17RA^low^ population, validating screen performance (Fig. 2A). The screen identified a broad network of genes with established roles in skin inflammation and immune regulation. Several top hits including ALOX5, PGF, LAMC1, and COL16A1 are involved in lipid metabolism and extracellular matrix organization, linking inflammatory signaling to structural remodeling in psoriatic skin. Moreover, the identification of chromatin regulators such as KAT5 and ASH2L suggests that IL17RA expression may be governed by epigenetic mechanisms in keratinocytes, consistent with their established roles in inflammatory gene regulation^14,15^. We also identified LCE3B, a well-established psoriasis susceptibility gene, as a hit^16^.

**Fig. 2.**
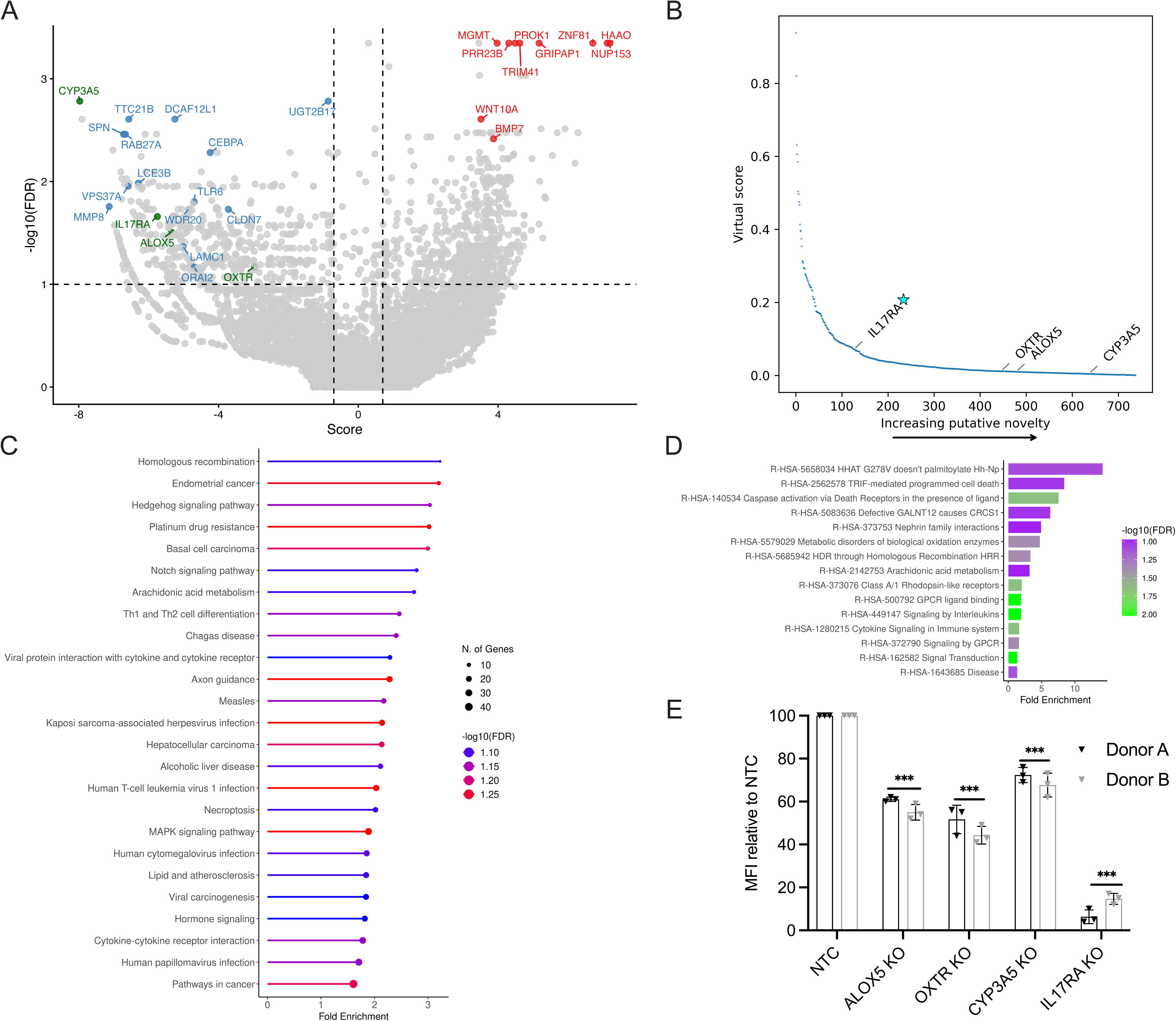
| Genome-wide CRISPR screen integrated with AI prioritization identifies regulators of IL17RA expression in primary human keratinocytes. **A**, Genome-wide screen results. Primary human adult epidermal keratinocytes from two independent donors were transduced with the Brunello sgRNA library and sorted for IL17RA surface expression. Plot shows log fold change (x-axis) versus −log10(FDR) (y-axis) for all genes. Genes causing decreased IL17RA surface expression are shown in blue (enriched in IL17RA^low^ population); those increasing IL17RA are in red (enriched in IL17RA^high^ population); those identified as novel by AI model as novel are in dark green. Dashed lines indicate significance thresholds (FDR < 0.1, |log fold change| > 0.6, ≥2 sgRNAs per gene). IL17RA was a positive control. **B**, AI-based novelty prediction. VirtualCRISPR model predictions for all screened genes, plotted by predicted hit probability (y-axis, virtual score) versus novelty rank (x-axis). Experimental screen hits with low predicted probabilities (below the curve) represent high-novelty, non-canonical regulators. OXTR, ALOX5, and CYP3A5 were prioritized as novel hits with lower predicted probability than IL17RA itself. **C**, KEGG pathway enrichment analysis of genes whose knockout decreased IL17RA expression. **D**, Reactome pathway enrichment analysis showing enrichment for GPCR signaling, signal transduction, and arachidonic acid metabolism among IL17RA regulators. **E**, Validation of top hits by targeted knockout in primary keratinocytes. Flow cytometric quantification of IL17RA surface expression in primary human adult epidermal keratinocytes from two independent donors following CRISPR-Cas9-mediated knockout of ALOX5, OXTR, CYP3A5, or IL17RA (two independent sgRNAs per gene). IL17RA mean fluorescence intensity (MFI) normalized to non-targeting control (NTC). Data are mean ± SEM (n = 2 biological donors, 3 technical replicates per donor per condition). ***P < 0.001, **P < 0.01, *P < 0.05 by two-way ANOVA versus NTC.

While these hits confirm the biological fidelity of the screen, existing psoriasis therapeutics already target well-characterized nodes in the IL-17 axis, meaning the highest clinical value lies not in rediscovering known regulators, but in identifying genes with strong experimental evidence and no prior association with IL17RA regulation in keratinocytes. To systematically prioritize such candidates, we employed the VirtualCRISPR model^12^, trained on the comprehensive BioGRID-ORCS database of functional genomic screens, which predicts the probability that disruption of a given gene in a specified cell type will produce a defined phenotype, addressing the question: “Does {perturbation} of {gene} in {cell type} result in {phenotype}?” (Fig. 1B). Critically, we deployed VirtualCRISPR not as a hit ranker but as a novelty filter. A high model-predicted probability signals that analogous gene-phenotype relationships already exist in the literature or training data; in other words, the biology is established. Conversely, experimental hits assigned low predicted probabilities represent genes that the model, drawing on the full landscape of published CRISPR screens, had no basis to predict, indicating genuinely uncharacterized regulatory mechanisms. This logic defines two distinct quadrants among enriched screen hits (Fig. 1C): genes with high experimental enrichment and high AI probability correspond to known regulators that serve as internal validation controls, while genes with high experimental enrichment and low AI probability constitute high-novelty candidates with unexplored biology. Importantly, this framework does not restrict or reduce the power of the unbiased genome-wide screen since all 19,000 screened genes and all experimental hits are fully retained. Instead, VirtualCRISPR annotates which hits represent new biology versus re-identification of known pathways, functioning as an orthogonal layer of prioritization rather than a filter on the experimental data.

We defined novel hits as genes falling in the high-novelty quadrant (Fig. 1C). Using this integrated framework, we identified Oxytocin receptor (OXTR), Arachidonate 5-lipoxygenase (ALOX5), and Cytochrome P450 3A5 (CYP3A5) as high-priority novel hits, each exhibiting a predicted probability lower than IL17RA knockout itself (Fig. 2B), confirming the model assigns no strong prior expectation to their role in IL17RA regulation in keratinocytes. OXTR and ALOX5 were prioritized given their novelty in the psoriasis context and tractable pharmacology with existing inhibitors, such as Cligosiban (OXTR) and Zileuton (ALOX5), enabling accelerated preclinical translation without requiring de novo drug development. CYP3A5 was included as a third validation candidate representing a mechanistically distinct xenobiotic metabolism axis.

Together, these candidates span neuroendocrine signaling (OXTR)^19^, lipid metabolism (ALOX5)^20^, and xenobiotic processing (CYP3A5)^21^, maximally diversifying the mechanistic insights extractable from the screen.

Pathway enrichment analysis showed arachidonic acid metabolism, interleukins signaling, and Notch signaling as enriched pathways (Fig. 2C-D). Notably, multiple hits converged on enzymatic nodes controlling arachidonic acid metabolism and eicosanoid synthesis, including, PLA2G2E, ALOX5, PTGES3, and PTGR2 (Supplementary Fig. 5). Cytokine-cytokine receptor interaction and signal transduction pathways were also enriched.

Beyond the arachidonic acid cascade, the screen identified innate immune pattern recognition as a critical node controlling IL17RA expression in keratinocytes. TLR6, a cell-surface Toll-like receptor that detects microbial components, and SARM1, a TIR-domain–containing adaptor that modulates Toll-like receptor signaling, emerged as regulators of IL17RA levels^17,18^. Analysis of GTExTCGA datasets revealed a strong positive correlation between ALOX5 and IL17RA expression (r = 0.89) across skin and spleen tissues, highlighting a robust transcriptional association between these genes (Supplementary Fig. 6). In addition, OXTR and CYP3A5 showed moderate correlation (r = 0.4 and 0.42) (Supplementary Fig. 7-8).

To validate the screen and assess the functional relevance of identified regulators, we performed targeted validation of the three prioritized candidates, OXTR, ALOX5, and CYP3A5. Each represents a distinct mechanistic axis implicated in epithelial immune regulation, spanning neuroendocrine signaling (OXTR)^19^, lipid metabolism (ALOX5)^20^, and xenobiotic processing (CYP3A5)^21^. Using CRISPR-Cas9 knockout with two sgRNAs per gene, we perturbed each locus in primary human adult epidermal keratinocytes derived from two independent donors to control for interindividual variability. Across all replicates and sgRNA pairs, genetic ablation of these genes significantly reduced IL17RA surface abundance, as quantified by flow cytometry (Fig. 2E), validating these genes as bona fide regulators of IL17RA expression in primary human keratinocytes.

We performed exploratory single cell RNA sequencing (scRNAseq) of peripheral blood mononuclear cells (PBMCs) from two psoriasis patients and two healthy controls. We generated 43,581 high-quality transcriptomes (mean ≈10,895 cells per sample) (Fig. 3A). We visualized the data using uniform manifold approximation and projection (UMAP), which delineated 21 transcriptionally distinct clusters corresponding to major immune cell lineages (Fig. 3B-C).

**Fig. 3.**
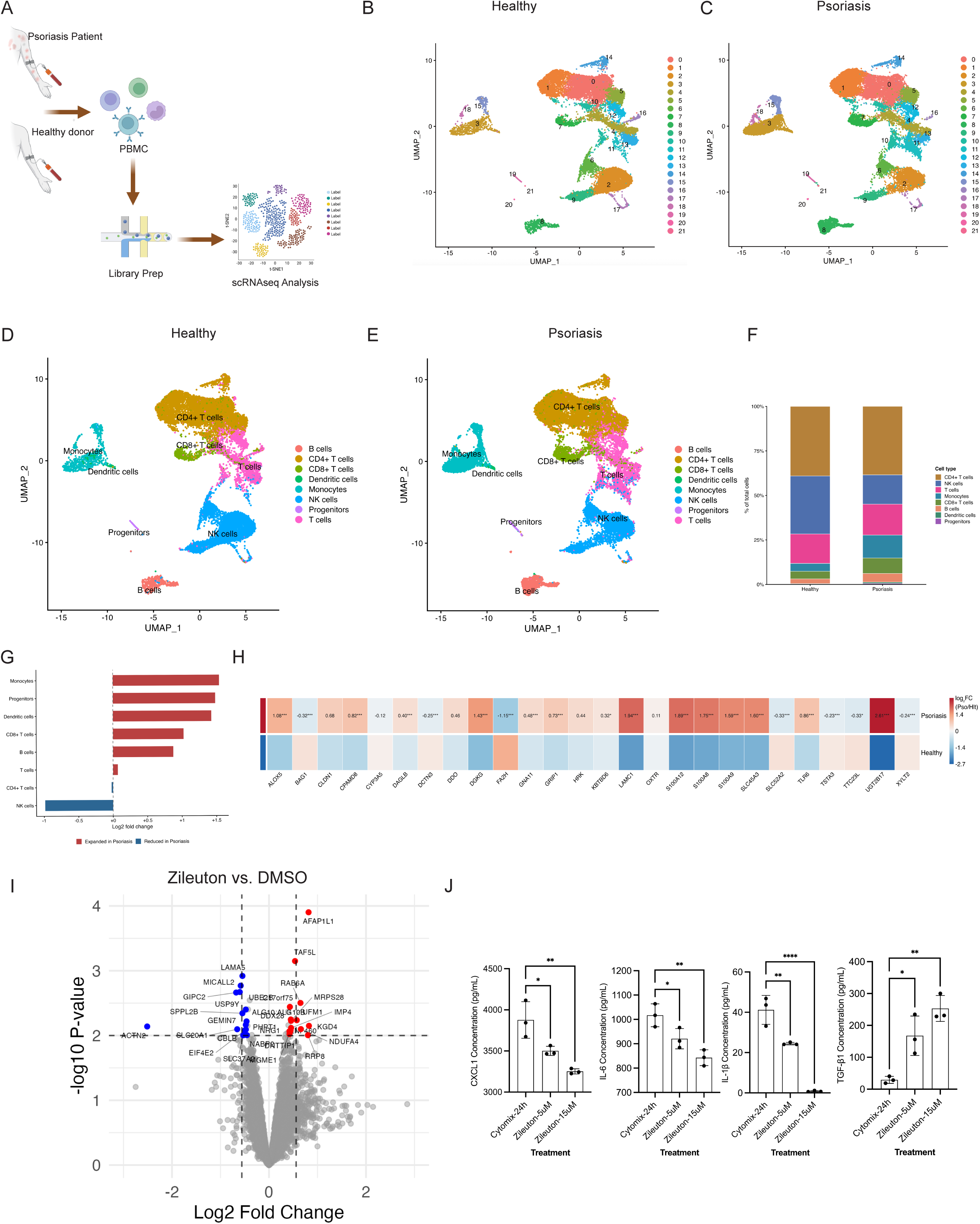
| Exploratory single-cell transcriptomic profiling reveals systemic expression of CRISPR screen hits, and Zileuton directly reprograms keratinocyte metabolism in human organotypic skin cultures. **A**, Experimental workflow for scRNA-seq of PBMCs using the 10X Genomics Chromium platform. **B**–**E**, Uniform Manifold Approximation and Projection (UMAP) visualization of 43,581 high-quality single-cell transcriptomes (mean 10,895 cells per sample) colored by Seurat clusters (**B**, healthy donors; **C**, psoriasis patients) or annotated cell types (**D**, healthy donors; **E**, psoriasis patients). Cell type annotation based on canonical markers: CD14^+^ monocytes (CD14, LYZ), CD4^+^ T cells (CD3E, CD4), CD8^+^ T cells (CD3E, CD8A), CD56^+^ NK cells (NCAM1), CD20^+^ B cells (MS4A1), and dendritic cells (CD1C, CLEC9A), and CD34⁺ progenitors. **F**, Percentage of different cell types in the healthy donors and psoriasis patients. **G**, Log2 fold change of different immune cell types in healthy donors versus psoriasis patients. **H**, Heatmap of 26 CRISPR screen hits (FDR ≤ 0.1 from primary keratinocyte screen, Fig. 2A) across PBMC immune cell populations. Color intensity indicates scaled average expression (z-score normalized within each gene). Wilcoxon rank-sum test with FDR correction, adjusted P < 0.05). **I**, Volcano plot of quantitative proteomic analysis (LC-MS/MS) from three-dimensional organotypic human skin raft cultures treated with Zileuton (15 μM) or DMSO vehicle for 48 h after 24 h cytomix stimulation. X-axis: log fold change (Zileuton/DMSO); y-axis: −log adjusted P value. Significantly downregulated proteins (P < 0.05) are shown in blue; upregulated proteins (P < 0.05) are shown in red. n = 4 independent organotypic cultures. **J**, ELISA quantification of the indicated secreted cytokines in culture supernatants from organotypic skin raft cultures treated with Zileuton or DMSO for 48 h after 24 h cytomix stimulation. Data are mean ± SEM (n = 3 independent organotypic cultures). *P < 0.05, **P < 0.01, ****P < 0.0001 by one-way ANOVA versus cytomix control

Annotation based on canonical marker genes identified CD14⁺ and FCGR3A⁺ (CD16⁺) monocytes; CD3E⁺CD4⁺ helper T cells; CD3E⁺CD8A⁺ cytotoxic T cells; NCAM1⁺ (CD56⁺) natural killer (NK) cells; MS4A1⁺ (CD20⁺) B cells; and CD1C⁺CLEC9A⁺ dendritic cells and CD34⁺ progenitors subsets (Fig. 3D-E). Psoriatic PBMCs showed increased frequency of monocytes, dendritic cells and CD8⁺ cells, while NK cells were reduced in psoriatic samples (Fig. 3F, G), in agreement with prior reports^22^.

We examined the differential expression of screen hits in psoriasis PBMCs. A plethora of screen hits showed elevated expression levels in psoriasis patients’ PBMC, including ALOX5, LAMC1, SLC45A3, and UGT2B17 which were elevated in psoriasis patients (Fig. 3H). Moreover, Psoriatic PBMCs exhibited elevated expression of alarmins (S100A8, S100A9, S100A12) (Fig. 3H), supporting the inflammatory phenotype of these samples^23^.

Given that ALOX5 regulates IL17RA (Fig. 2E) and is elevated in psoriasis patients PBMCs (Fig. 3H), these findings suggest ALOX5 may participate in the psoriasis-associated inflammatory process in both keratinocytes and circulating immune cells, although the functional relationship between these compartments remains to be determined. Furthermore, the convergence of multiple hits on arachidonic acid metabolism, together with the availability of Zileuton as an FDA-approved ALOX5 inhibitor, we prioritized ALOX5 for therapeutic validation.

This selection will enable accelerated progression to preclinical models without requiring a new drug development pipeline. A critical question remained: does Zileuton act directly on keratinocytes, or does it require immune cell intermediates? To address this, we turned to 3D organotypic human skin culture which is a physiologically relevant human tissue model that recapitulates the stratified architecture, calcium-dependent differentiation gradients, and barrier function of native epidermis. This system isolates primary keratinocyte responses from confounding immune signals, providing unambiguous evidence for cell-autonomous drug effects. We established organotypic cultures from HEKa, treated them with Zileuton (15 μM) for 2 days, and performed quantitative proteomic profiling.

The results were striking, Zileuton induced profound metabolic reprogramming of human keratinocytes in the absence of immune cells (Fig. 3I), demonstrating the therapeutic mechanism operates directly at the epithelial level rather than through secondary immunomodulation.

At the molecular level, Zileuton treatment was associated with coordinated suppression of three pathogenic programs. First, interferon-stimulated genes (OAS2, OAS3, IFITM3, UBE2L6) were markedly downregulated (Supplementary table 6), consistent with attenuation of keratinocyte-intrinsic type I interferon signaling, which has been implicated in psoriatic inflammation particularly in early disease and interferon-driven subtypes. Second, cholesterol biosynthesis enzymes (HMGCS1, IDI1) were reduced (Supplementary table 6), suggesting decreased mevalonate pathway activity. Third, amino acid transporters essential for anabolic growth and proliferation (SLC7A5, SLC7A11, SLC1A5) were markedly suppressed (Supplementary table 6), consistent with reduced proliferative drive^24^.

Concomitantly, Zileuton treatment drove a robust metabolic reprogramming. Keratinocytes showed a marked induction of mitochondrial biogenesis machinery (TFAM; multiple mitochondrial ribosomal, and respiratory chain proteins, including NDUFA13, NDUFS6, NDUFV2) (Supplementary table 6). This mitochondrial expansion supported coordinate activation of fatty-acid β-oxidation (HADHA, HADHB, ACAA2, DECR1, DECR2, ECI2) and ketone-body utilization (OXCT1) (Supplementary table 6), reflecting a fundamental metabolic shift from anabolic lipid synthesis, which fuels keratinocyte hyperproliferation in psoriasis^25^, toward catabolic oxidative metabolism, consistent with the metabolic phenotype of terminally differentiated keratinocytes.

Together, findings from the 3D organotypic model support three key principles: (1) Zileuton exerts direct effects on keratinocytes independent of immune cell contributions; (2) these effects are preserved within stratified epithelium, supporting the feasibility of topical delivery strategies; and (3) Zileuton promotes a metabolic and transcriptional state aligned with epidermal homeostasis rather than psoriatic hyperproliferation.

Critically, these changes were not isolated effects but rather a coordinated transcriptional program characterized by simultaneous suppression of pathogenic inflammatory and proliferative pathways coupled with induction of differentiation and oxidative metabolism, indicating that Zileuton triggers a fundamental cell fate transition.

Additionally, Zileuton treatment increased TGF-β1 while suppressing pro-inflammatory cytokines CXCL1, IL-6, and IL-1β secretions shifting the balance toward immune-regulatory signaling (Fig. 3J). Collectively, these findings indicate that Zileuton reprograms keratinocytes by suppressing inflammatory and proliferative networks while promoting mitochondrial metabolism and epithelial homeostasis, explaining its therapeutic efficacy through direct epithelial effects.

To evaluate the therapeutic potential of OXTR and ALOX5 inhibition in vivo, we formulated Zileuton and Cligosiban as topical gels using Carbomer 940 as a delivery matrix^26^. Carbomer 940 was selected for its rapid absorption, ease of application, and lack of inherent anti-inflammatory activity (Supplementary Fig. 9A-D).

We employed the imiquimod (IMQ)-induced psoriasis model, a well-validated preclinical murine model for studying IL-17-driven skin inflammation^27^. Mice received Zileuton gel, Cligosiban gel, anti-IL17RA antibody, vehicle gel, or IMQ only. The topical IMQ (5% Aldara cream) control group was not treated with a therapeutic intervention (Fig. 4A). We monitored disease severity daily using PASI scoring and harvested skin on Day 7 for flow cytometric immunophenotyping and proteomic analysis.

**Fig. 4.**
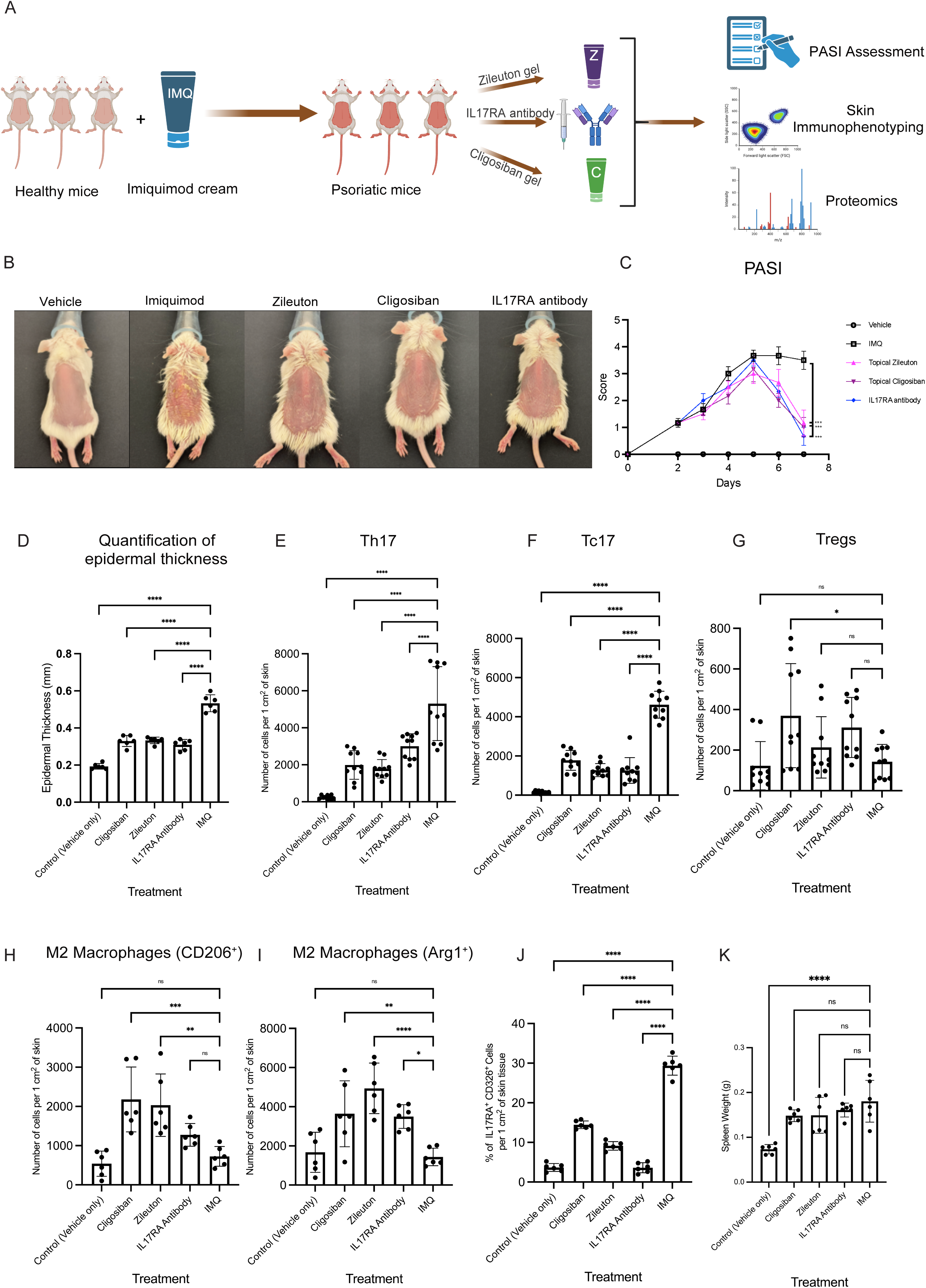
| Topical ALOX5 and OXTR inhibition suppresses psoriasiform inflammation in the imiquimod mouse model. **A**, Experimental design. Psoriasiform dermatitis was induced in female BALB/c mice (6–8 weeks) by daily topical application of imiquimod (IMQ; 5% Aldara cream, 62.5 mg) to dorsal skin for 7 consecutive days. Topical Zileuton gel and topical Cligosiban gel were applied on day 3 to 6, systemic anti-IL17RA monoclonal antibody, vehicle gel (Carbomer 940, without IMQ), or no treatment control (IMQ + vehicle gel). Disease severity was assessed daily using the modified Psoriasis Area and Severity Index (PASI; cumulative score of erythema, scaling, and thickness). On day 7, mice were euthanized, and dorsal skin was harvested for flow cytometric immunophenotyping, histological analysis, and proteomic profiling. n = 6 mice per group. **B**, Representative macroscopic images of dorsal skin on day 7 showing reduced erythema, scaling, and thickening in Zileuton-, Cligosiban-, and anti-IL17RA antibody-treated mice compared to IMQ-only control. **C**, Daily PASI scores over 7 days. Both topical treatments (Zileuton and Cligosiban) significantly reduced disease severity with kinetics and magnitude comparable to systemic IL-17RA blockade. Data are mean ± SEM. ***P < 0.001 for Zileuton, Cligosiban, and anti-IL17RA antibody versus IMQ-only by two-way repeated measures ANOVA with Tukey’s multiple comparisons test. **D**, Quantification of epidermal thickness. Data are mean ± SEM from ≥5 measurements per mouse. ****P < 0.0001, ***P < 0.001 by one-way ANOVA with Tukey’s multiple comparisons test versus IMQ-only control. **E–G**, Flow cytometric quantification of skin-infiltrating T cell subsets (gating strategy in Supplementary Fig. 13). **E**, Th17 cells (CD45^+^CD3^+^CD4^+^RORγt^+^IL-17A^+^). **F**, Tc17 cells (CD45^+^CD3^+^CD8^+^RORγt^+^IL-17A^+^). **G**, Tregs (CD45^+^CD4^+^CD25^+^Foxp3^+^). Cell numbers normalized to 1 cm^2^ of skin tissue. **H**, CD206^+^ macrophages. **I**, Arg1^+^ macrophages. Topical treatments (Zileuton and Cligosiban) uniquely increased M2 polarization, an effect not observed with systemic anti-IL17RA antibodies. **J**, Percentage of IL17RA^+^ cells among total skin cells by flow cytometry, demonstrating receptor downregulation following topical treatments. **K**, Spleen weight as a measure of systemic immune activation. Data in D–K are mean ± SEM. ****P < 0.0001, ***P < 0.001, **P < 0.01, *P < 0.05, ns (not significant) by one-way ANOVA with Tukey’s multiple comparisons test versus IMQ-only control unless otherwise indicated.

Quantitative assessment using the Psoriasis Area and Severity Index (PASI) showed that topical OXTR or ALOX5 inhibition substantially reduced IMQ-induced inflammation (Fig. 4B). Cligosiban and Zileuton significantly reduced cumulative PASI scores relative to IMQ-only controls, with clinical improvement evident by Day 5 and sustained to almost full resolution by Day 7 (Fig. 4C). Both compounds achieved efficacy comparable to systemic anti-IL17RA antibody, establishing that topical targeting can recapitulate systemic blockade.

Histomorphometric quantification of epidermal thickness revealed significant normalization of acanthosis in both topical treatment groups (Fig. 4D). The magnitude of epidermal resolution was comparable to that observed with antibody treatment, reflecting effective suppression of keratinocyte hyperproliferation and IL-17-driven pathology.

To investigate the immunological basis of disease improvement, we quantified immune cell infiltration in the skin by flow cytometry. Topical treatments reduced Th17 cell frequencies as effectively as systemic antibody treatment (Fig. 4E). Both topical inhibitors significantly reduced Th17 and Tc17 cell populations (Fig. 4F). Additionally, Cligosiban and anti-IL17RA antibody treatment increased CD4⁺ regulatory T-cells (Tregs) frequencies, whereas Zileuton showed no increase in Tregs frequencies (Fig. 4G).

Analysis of the myeloid compartment revealed a pronounced shift toward an anti-inflammatory, M2-like phenotype. Notably, CD206⁺ and Arg1⁺ M2 macrophages were significantly enriched following topical treatment with Cligosiban or Zileuton, whereas systemic IL17RA antibody administration selectively increased Arg1⁺ M2 macrophages (Fig. 4H–I). Consistent with in vitro observations, IL17RA⁺ epidermal cells were markedly reduced in mice receiving either topical treatment compared to IMQ-only controls, confirming that both compounds downregulate receptor expression in vivo (Fig. 4J). Evaluation of systemic inflammatory burden through spleen weight measurements showed that IMQ induced splenomegaly across all treatment groups relative to healthy controls, indicating sustained systemic immune activation (Fig. 4K)^28^.

Finally, Ki67 staining demonstrated that all treatments effectively normalized epidermal proliferation to baseline levels (Supplementary Fig. 10), collectively supporting restoration of tissue homeostasis.

Together, these results show that topical administration of Cligosiban and Zileuton effectively suppresses psoriatic inflammation through multiple mechanisms: reducing IL17RA expression, suppressing pathogenic Th17/Tc17 responses, and polarizing macrophages toward an anti-inflammatory phenotype. The multi-pronged immunomodulation observed establishes proof-of-concept for topical small-molecule therapy targeting keratinocyte-specific pathways in psoriasis.

To understand global proteomic changes induced by each treatment, we analyzed dorsal skin tissue. Examining the proteins with significant differential abundance across treatments provides mechanistic granularity to the global proteomic trends. Anti-IL17RA antibody treatment in mice suppressed a multi-layered effector program encompassing neutrophil-derived antimicrobial proteases (Fig. 5A, Supplementary table 7) (Elane, Ctsg, Prtn3, Mpo) and alarmins (S100a8, S100a9, Lcn2), that collectively drive psoriasiform inflammation^29,30^. The coordinate downregulation of keratinocyte-derived antimicrobial peptides (Camp, Defb3) and neutrophil-associated granule constituents (Ltf, Pglyrp1) supports attenuation of the IL-17–driven keratinocyte–neutrophil inflammatory loop at both epithelial and myeloid levels, consistent with effective on-target pathway suppression^31^.

**Fig. 5.**
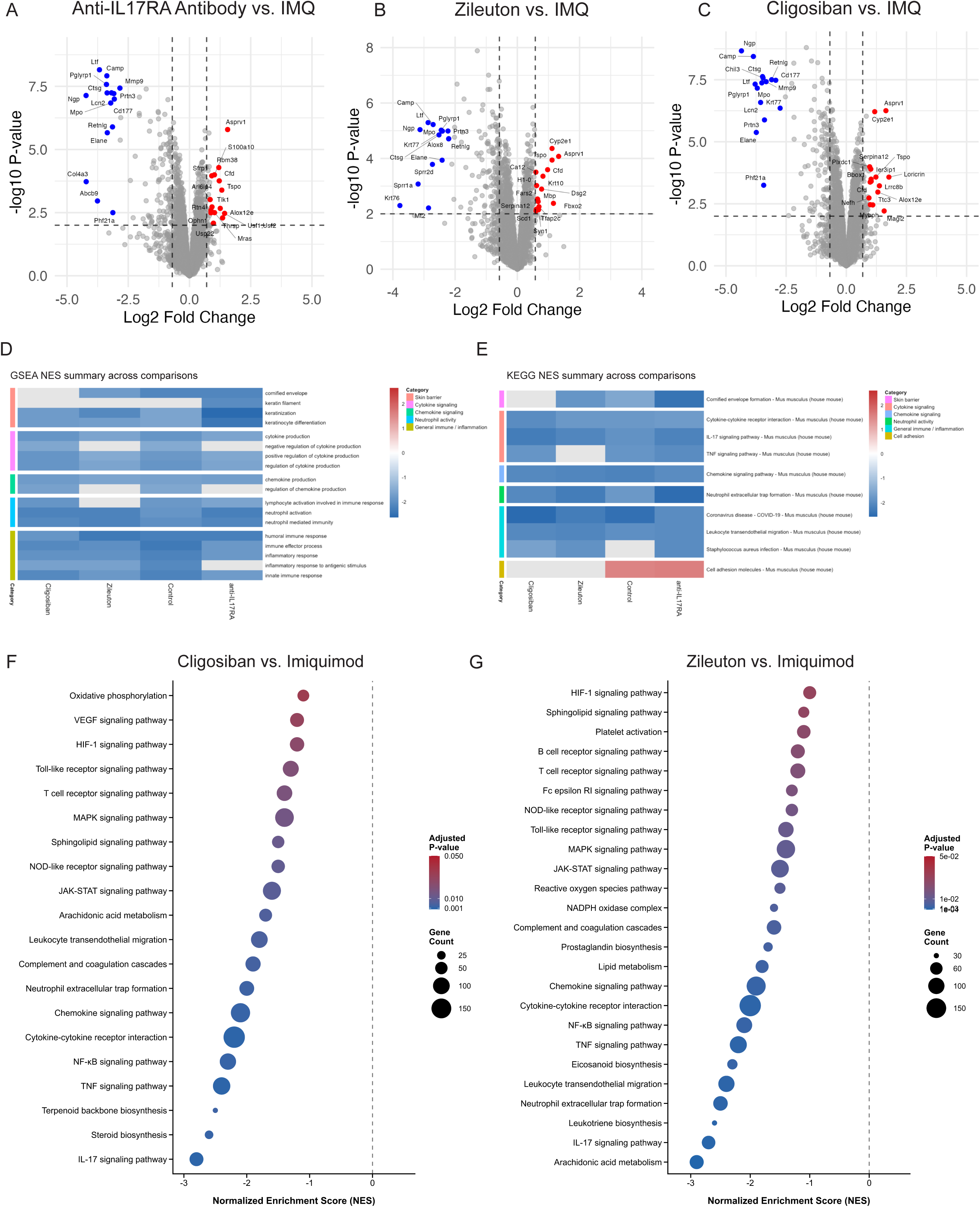
| Proteomic profiling reveals convergent suppression of inflammatory effectors and distinct metabolic reprogramming by systemic IL17RA blockade and topical ALOX5 or OXTR inhibition. Quantitative proteomic analysis of dorsal skin tissue from imiquimod (IMQ)-treated mice (day 7) following therapeutic intervention. **A**–**C**, Volcano plots of differential protein expression comparing **A**, anti-IL17RA antibody; **B**, Zileuton (topical ALOX5 inhibitor); or **C**, Cligosiban (topical OXTR antagonist) versus IMQ-only control. Proteins were extracted from skin tissue and analyzed by label-free quantitative LC-MS/MS (n = 6 mice per group, replicates per biological sample). Each point represents one protein. X-axis: log fold change; y-axis: −log P value. Blue points: significantly downregulated proteins (log fold change < −0.5, P < 0.05); red points: significantly upregulated proteins (log fold change > 0.5, P < 0.05); gray points: not significant. Dashed lines indicate significance thresholds. Statistical significance determined by two-sample t-test with Benjamini–Hochberg false discovery rate (FDR) correction. **D**,**E**, Pathway enrichment analysis across all treatment comparisons using significantly altered proteins (|log fold change| > 0.5, adjusted P < 0.05). **D**, Gene Set Enrichment Analysis (GSEA) of MSigDB Hallmark gene sets showing normalized enrichment scores (NES). Negative NES indicates pathway downregulation. **E**, KEGG pathway enrichment scores. **F**-**G**, Comprehensive KEGG pathway enrichment profiles for **F**, Cligosiban versus IMQ **G**, Zileuton versus IMQ. Pathways are ranked by NES. Color intensity indicates adjusted P-value.

Matrix metalloproteinases (Mmp8, Mmp9) and neutrophil-specific markers (Ngp, Cd177) were similarly suppressed, indicating reduced tissue remodeling and neutrophil infiltration.

Collectively, these proteomic signatures are consistent with the established role of IL-17RA signaling in coordinating keratinocyte activation and neutrophil recruitment. Furthermore, the upregulation of late differentiation markers (Ivl, Sbsn, Scel), together with desmosomal adhesion proteins (Dsg1a) and extracellular matrix components (Fbln1, Fmod, Serpinf1), indicates improved intercellular cohesion and barrier restoration.

Zileuton treatment recapitulated many of the anti-inflammatory effects observed with IL-17RA blockade (Fig. 5B, Supplementary table 8) including suppression of neutrophil effectors (Elane, Ctsg, Prtn3, Mpo, Cd177), antimicrobial peptides and alarmins (Camp, Defb3, S100a8, S100a9), matrix metalloproteinases (Mmp8, Mmp9). Directly validating on-target pharmacology, Zileuton treatment reduced the abundance of most components of the leukotriene biosynthetic pathway in psoriatic skin, including Alox5, Alox5ap, Pla2g4b, Pla2g4e, and Lta4h. Consistent with reduced oxidative effector capacity, Zileuton decreased components of the NADPH-oxidase complex (Ncf1, Ncf2, Ncf4).

To validate this circuit mechanistically, we interrogated the relationship between ALOX5 and IL-17RA in HEKa. ALOX5 knockout resulted in profound IL-17RA downregulation compared to WT cells (Supplementary Fig. 11A), while exogenous Leukotriene B4 (LTB4), a downstream leukotriene mediator of ALOX5 activity, dose-dependently increased IL17RA surface expression (Supplementary Fig. 11B). Surface flow cytometry confirmed that Zileuton treatment significantly reduced IL17RA surface levels (Supplementary Fig. 11). Notably, this reduction was fully rescued by Dynasore, a dynamin inhibitor that blocks clathrin-mediated endocytosis, indicating that leukotriene signaling promotes IL-17RA surface stability in a dynamin-dependent manner.

Together, these findings support a model in which leukotriene activity sustains IL-17RA surface availability by limiting receptor internalization. (Supplementary Fig. 11C).

Proteomic analyses revealed concurrent upregulation of trafficking components implicated in endocytic and degradative routing, including the actin motor myosin VI (Myo6), the lysosomal positioning factor (Rab34), and the sorting nexin (Snx7)^32,33^. Concurrent suppression of vesicular trafficking regulators involved in endosomal sorting, lysosome-related organelle biogenesis, and secretory pathway organization (Rab27a, Rab32, Trappc3) suggests altered membrane trafficking dynamics that may reduce receptor surface replenishment through multiple routes. Together with the demonstrated requirement for dynamin-dependent internalization, this coordinated trafficking signature is consistent with reduced receptor recycling and altered post-endocytic routing that diminishes surface availability under Zileuton treatment.

Beyond the shared neutrophil-keratinocyte axis, Zileuton selectively reduced acute phase reactants and complement cascade components (C3, Cfb, Hp, Itih2, Itih4, Lbp) mirroring the anti-inflammatory signature observed with IL-17RA antibody-treated cohort as well as Hmgcs1 and IDI1 which were also downregulated in the 3D organotypic culture. The in vivo validation bridges the gap between our mechanistic in vitro findings and 3D organotypic validation, supporting the concept that keratinocyte-directed small molecules can modulate epithelial and inflammatory programs in psoriatic skin.

Cligosiban treatment elicited a proteomic response that phenocopied IL-17RA blockade on core psoriasiform effectors (Fig. 5C, Supplementary table 9), including neutrophil proteases (Elane, Ctsg, Prtn3, Mpo), antimicrobial peptides and alarmins (Camp, Defb3, S100a8, S100a9, Lcn2), and hyperproliferative keratinocyte markers (Sprr2d, Sprr2f) yet established a distinctive anti-inflammatory axis. Like Zileuton and IL-17RA blockade, Cligosiban suppressed multiple core components of the NADPH oxidase complex (Ncf1, Ncf2, Ncf4, Cyba, Cybb) and leukotriene biosynthetic enzymes (Alox5, Alox5ap, Pla2g4b, Pla2g4d, Pla2g4e), showing shared control over oxidative burst and eicosanoid metabolism.

However, Cligosiban induced a far more comprehensive metabolic reprogramming signature than ALOX5 pathway inhibition. Cligosiban uniquely reduced abundance of multiple enzymes across the mevalonate–cholesterol biosynthesis pathway, including the rate-limiting enzyme HMG-CoA reductase (Hmgcr) alongside downstream sterol synthesis machinery (Hmgcs1, Mvd, Fdps, Fdft1, Sqle, Lss, Cyp51a1, Sc5d, Hsd17b7, Msmo1, Pmvk). Cligosiban also suppressed sphingolipid biosynthesis enzymes (Sptlc1, Sptlc2, Degs1, Cers3) and fatty acid elongases (Elovl1). The coordinate regulation suggests that OXTR signaling modulates membrane lipid biosynthetic programs that sustains inflammatory keratinocyte hyperproliferation and barrier dysfunction^34,35^.

Indeed, OXTR knockout in HEKa cells resulted in IL-17RA downregulation alongside reduced calmodulin levels (Supplementary Fig. 12A), and Oxytocin treatment increased IL-17RA surface expression. However, the effects were abolished by calcium chelation (BAPTA-AM) or calcineurin inhibition (tacrolimus) (Supplementary Fig. 12B-E). These findings suggest that OXTR-mediated calcium mobilization regulates IL-17RA expression in a calcineurin-dependent manner, consistent with involvement of the calmodulin–calcineurin–NFAT signaling axis^36^.

Supporting this mechanism, the Cligosiban in vivo proteomic signature revealed coordinated suppression of calmodulin effectors (Marcks), calcium-activated enzymes (Pla2g4b, Pla2g4d, Pla2g4e, Padi4), and AP-1 components (Fosl2, Junb), suggesting attenuation of calcium-dependent inflammatory signaling pathways (Fig. 5C, Supplementary table 9). Marcks, a direct calmodulin substrate, suppressed alongside calcium-sensing protein Efhd2, corroborating attenuation of OXTR-dependent intracellular calcium signaling following receptor antagonism.

Cligosiban also reduced multiple inflammatory mediators and AP-1 transcription factor components (Fosl2, Junb), which are known to contribute to inflammatory gene expression programs. Reduced abundance of Il1a, Ptges, Cd44, and Osmr further indicates attenuation of pathways involved in inflammatory amplification, prostaglandin synthesis, cell adhesion, and cytokine signaling^37^.

Beyond metabolic and transcriptional regulation, Cligosiban was associated with reduced abundance of innate immune and danger-sensing pathway components. The coordinate downregulation of Sting1 (cGAS-STING pathway)^38^, complement anaphylatoxin receptor C5ar1, formyl peptide receptor Fpr2^39^, and the alarmin IL-36γ (Il36g) suggests that OXTR antagonism dampens danger signal sensing and inflammatory amplification loops.

The convergent suppression of neutrophil effectors, antimicrobial peptides, and inflammatory mediators across mechanistically distinct interventions, including direct IL-17RA neutralization, via ALOX5 and OXTR inhibition, reinforces IL-17RA as a central signaling node orchestrating psoriasiform inflammation and establishes OXTR and ALOX5 as pharmacologically tractable upstream regulators of IL-17RA driven pathology. However, proteomic profiles diverged beyond the shared IL-17RA dependent inflammatory axis, revealing treatment-specific metabolic and transcriptional reprogramming signatures that account for differential therapeutic efficacy.

Pathway enrichment analysis showed that all three treatments suppressed core inflammatory pathways: IL-17, TNF, cytokine-cytokine receptor, and chemokine signaling (Fig. 5D-E). The consistent downregulation of these modules across all treatments validates that each therapeutic approach effectively disrupts the keratinocyte-immune cell inflammatory amplification cascade.

Despite convergence on common inflammatory pathways, each experimental condition maintained unique mechanistic signatures (Fig. 5F-G). Zileuton showed the most pronounced suppression of arachidonic acid metabolism, consistent with ALOX5 inhibition and blockade of the leukotriene biosynthesis axis. In contrast, Cligosiban selectively downregulated steroid and terpenoid backbone biosynthesis pathways, including key rate-limiting enzymes in cholesterol synthesis, highlighting OXTR-dependent lipid metabolic reprogramming distinct from leukotriene-driven inflammation.

All three interventions converged on conserved epidermal resolution programs, characterized by coordinated barrier restoration and suppression of inflammatory effector pathway.^41,42^, with coordinated upregulation of the profilaggrin-processing protease Asprv1, consistent with partial restoration of keratinocyte terminal proteolytic homeostasis. However, the three interventions differed markedly in the breadth of terminal differentiation programs induced.

Zileuton upregulated the early suprabasal differentiation marker Krt10 and several adhesion and matrix-remodeling modules, including the desmosomal protein Dsg2, tight-junction scaffolds (Tjp3, Cgn, Cgnl1), the hemidesmosomal cytolinker Dystonin, and basement membrane and extracellular matrix components (Lama4, Fbln1, Fbln5), consistent with early reinforcement of epidermal structural integrity. IL-17RA blockade achieved intermediate differentiation restoration, upregulating a partial late-stage cornified envelope program (Ivl, Sbsn, Scel) together with shared extracellular matrix components (Fbln1) and the desmosomal adhesion protein Dsg1a, consistent with enhanced epidermal cohesion. Cligosiban elicited the broadest differentiation signature, strongly inducing terminal differentiation markers including Loricrin, Involucrin, Filaggrin, Corneodesmosin, Suprabasin, and Sciellin alongside differentiation keratins Krt10 and Krt1, suggesting more extensive reactivation of the stratified epidermal differentiation program.

The graded restoration of epidermal terminal differentiation across the three interventions from structural reinforcement under Zileuton, through partial cornified envelope induction under IL-17RA blockade, to broader terminal differentiation activation under Cligosiban, suggests that OXTR antagonism may modulate not only inflammatory signaling but also epithelial differentiation programs relevant to chronic psoriatic barrier dysfunction.

## Discussion

We report the first genome-wide CRISPR screen performed in primary human adult keratinocytes to identify regulators of IL17RA expression. We integrated unbiased functional genomics with AI-driven target prioritization and identified ALOX5 and OXTR as therapeutically tractable modulators of IL17RA expression. Topical delivery of their respective inhibitors, Zileuton and Cligosiban, achieved therapeutic effects comparable to systemic IL17RA blockade in preclinical models, providing proof-of-concept for keratinocyte-directed small molecule strategies targeting upstream regulators of IL-17 signaling.

ALOX5 emerged as a key regulator that appears to sustain IL17RA surface expression by limiting receptor internalization. Pharmacologic inhibition with Zileuton promoted IL17RA internalization in a dynamin-dependent manner while reducing neutrophil effectors, NADPH oxidase components, and markers associated with neutrophil extracellular trap formation. These coordinated effects recapitulated core inflammatory features of systemic IL17RA blockade through inhibition of a single enzymatic pathway. In line with its established role as an ALOX5 inhibitor and its FDA-approved safety profile in asthma, Zileuton may offer a feasible path toward topical clinical translation, particularly for localized disease or for patients who demonstrate inadequate responses to biologic therapy, although this requires prospective clinical evaluation^42^.

OXTR represents a less conventional discovery. As a Gq-coupled GPCR, OXTR regulates IL17RA expression through a calcineurin-dependent mechanism consistent with involvement of calcium–calcineurin–NFAT signaling. Beyond receptor regulation, the breadth of its proteomic footprint, encompassing mevalonate–cholesterol biosynthesis, sphingolipid metabolism, oxidative burst machinery, and innate immune sensing suggests that OXTR occupies a central regulatory position in keratinocyte inflammatory competence. Clinically approved OXTR antagonists, including Atosiban and Barusiban (currently used as tocolytics), may offer accelerated repurposing opportunities given their established human safety profiles, although dermatological formulation, dosing, and efficacy will require dedicated investigation^43^.

Our work establishes a rapid, integrated discovery platform combining primary cell CRISPR screening, AI-guided prioritization, and multi-omics validation to systematically identify candidate therapeutic targets in complex inflammatory diseases, advancing the path from mechanistic discovery to preclinical proof of concept.

The convergence of leukotriene and calcium signaling on IL17RA regulation reveals previously underappreciated mechanisms governing keratinocyte inflammatory competence and highlights novel intervention points in psoriasis. The therapeutic effects of topical small molecules targeting these pathways suggest the potential for alternatives to systemic biologics, with implications for improving treatment accessibility and reducing therapeutic burden. More broadly, this study provides a blueprint for how AI-enabled functional genomics can surface non-obvious therapeutic targets and accelerate translation from primary cell discovery to preclinical validation in complex inflammatory diseases.

## Author contributions

A.M.A., S.O.K., and A.A.K. conceived and designed the study. C.Z. and M.S. conducted most of the experiments and performed the data analyses. S.A. conducted the animal work with assistance from C.Z.. J.F., J.M.,and P.Y.L provided technical assistance in molecular biology experiments. A.L. performed the mass spectrometry experiments under the supervision of R.M.

S.S. and J.P. developed the large language models and performed the coding under A.A.K supervision.. A.T., A.K., and E.W. helped with the initial validation experiments. A.M.A. and S.O.K. supervised the project. A.M.A., S.O.K., M.S., and C.Z. wrote the manuscript with input from all authors.

## Acknowledgments

This work was supported by the Chan Zuckerberg Biohub Chicago. We thank the operations team at CZ Biohub Chicago for their support and assistance throughout this project. We also thank Hope Burks, Quinn Roth-Carter, and Hanadi Almazroue for helpful discussions and input on the project.

## Ethics Declaration

The authors have filed intellectual property related to the results reported in this study

## Notes

### Competing Interest Statement

The authors have declared no competing interest.

